# Interneuron-targeted disruption of *SYNGAP1* alters sensory representations in neocortex and impairs sensory learning

**DOI:** 10.1101/2022.09.27.509690

**Authors:** Meiling Zhao, Sung Eun Kwon

## Abstract

*SYNGAP1* haploinsufficiency in humans results in severe neurodevelopmental disorders characterized by intellectual disability, autism, epilepsy, and sensory processing deficits. However, circuit mechanisms that underlie *SYNGAP1*-related neurodevelopmental disorders are poorly understood. A decrease of SynGAP in mice causes cognitive and behavioral deficits in part by disrupting the development of excitatory glutamatergic connections. Whether and to what extent SynGAP functions in inhibitory circuits remain unclear. We show that interneuron-specific *SYNGAP1* heterozygous mice display learning deficits characterized by elevated behavioral responses in the absence of relevant sensory input and premature responses to a sensory input unrelated to reward acquisition. These behavioral deficits are associated with specific circuit abnormalities within primary somatosensory cortex, characterized by increased detrimental correlations and elevated responses to irrelevant sensory stimuli. Collectively, we show that a decrease of SynGAP in inhibitory interneurons disrupts sensory representation in the primary sensory cortex, which likely contributes to behavioral deficits.

## Introduction

*De novo* loss-of-function variants of the gene *SYNGAP1* cause neurodevelopmental disorders characterized by intellectual disability, developmental delay, autism, schizophrenia, and epilepsy (Berryer et al., 2013; De Rubeis et al., 2014; Mignot et al., 2016; Satterstrom et al., 2020). *SYNGAP1* encodes a synaptically localized GTPase-activating protein (SynGAP) that enhances the intrinsic GTPase activity of H-Ras and interacts with a synaptic scaffolding protein PSD-95 (Chen et al., 1998; Kim et al., 1998; Komiyama et al., 2002). The importance of SynGAP in neuronal maturation, synapse development and synaptic plasticity is well-documented both *in vitro* and *in vivo* (Araki et al., 2015; Barnes et al., 2015; Clement et al., 2012; Kim et al., 2003; Komiyama et al., 2002; Llamosas et al., 2021). However, it is unclear how pathogenic *SYNGAP1* variants impact neural circuits and lead to behavioral abnormalities.

Although *SYNGAP1* is predominantly expressed in excitatory neurons of forebrain structures including the cerebral cortex and hippocampus, its expression is also detected in the inhibitory neurons in the forebrain (Berryer et al., 2016; Su et al., 2019; Velmeshev et al., 2019; Zhang et al., 1999). The questions of whether and to what extent SynGAP functions in inhibitory circuits have received relatively less attention. SynGAP was shown to be essential for the migration of inhibitory neurons during development (Su et al., 2019). Pan-neuronal haploinsufficiency of *SYNGAP1* impacts inhibitory as well as excitatory neurons (Michaelson et al., 2018; Sullivan et al., 2020). Importantly, a selective loss of *SYNGAP1* in GABAergic neurons generated in the medial ganglionic eminence (MGE) disrupts the ability of parvalbumin-expressing (PV) inhibitory cortical interneurons to provide perisomatic inhibition in a cell-autonomous manner (Berryer et al., 2016). The resulting loss of inhibition onto excitatory pyramidal neurons may contribute to altered cortical gamma oscillations, cognitive deficits, and impaired social interaction (Berryer et al., 2016). However, circuit-level consequences of reducing SynGAP in GABAergic neurons remain unclear, since neural and behavioral phenotypes have not been analyzed in the same animals.

GABAergic neurons are critically important for the regulation of cortical activity and are frequently disrupted in neurodevelopmental disorders (Contractor et al., 2021; Isaacson and Scanziani, 2011; Velmeshev et al., 2019). Dysfunctional inhibitory circuits can negatively impact sensory processing and learning due to altered sensory tuning, increased stimulus sensitivity, or aberrant neural correlations in the network without concomitant increases in spiking (Chen et al., 2020; Goel et al., 2018; Goncalves et al., 2013). One way inhibitory neurons modulate network activity is by imposing a strict ‘window of opportunity’ for temporal integration of synaptic inputs such that relevant sensory information is represented in the brain while irrelevant inputs are filtered out (Pouille and Scanziani, 2001). The potential therapeutic efficacy of targeting inhibitory neurons has been demonstrated in mouse models of autism including Fragile-X syndrome (Goel et al., 2018). Addressing whether and how *SYNGAP1* haploinsufficiency in inhibitory neurons causes circuit-level defects is important for basic understanding and potential therapeutic intervention of cognitive difficulties in *SYNGAP1*-related neurodevelopmental disorders.

To investigate how inhibitory interneuron-specific *SYNGAP1* haploinsufficiency impacts cortical circuit and sensory perception, we generated a haploinsufficient mouse model in which a copy of *SYNGAP1* gene was selectively knocked out in vesicular GABA transporter (Vgat)-expressing neurons by using Cre-lox. We then probed circuit-level alterations using two-photon calcium imaging of the whisker primary somatosensory cortex (wS1), in order to capitalize on its well-defined circuitry. This approach was similar to prior studies, which established the link between *SYNGAP1* haploinsufficiency and abnormal whisker input processing (Michaelson et al., 2018). Mice were head-restrained and trained to report the detection of whisker vibration by licking a reward port. Each trial began with a brief auditory tone indicating trial initiation. We monitored neural responses in layer 2/3 (L2/3) of wS1 while mice performed this task. These experiments allowed us to assess both behavioral and circuit-level consequences of inhibitory interneuron-specific *SYNGAP1* haploinsufficiency in the same animal.

We show that interneuron-specific *SYNGAP1* heterozygous mice (Vgat-Het) display learning deficits characterized by elevated behavioral responses in the absence of relevant sensory input (whisker vibration) and premature responses to a sensory input unrelated to reward acquisition (auditory tone). These behavioral deficits are associated with specific circuit abnormalities within wS1. Pairwise noise correlations are widely distributed with greater variability in Vgat-Het mice. The average noise correlation was slightly yet significantly elevated in trained Vgat-Het mice. Abolishing noise correlations improved decoding of stimulus identity from L2/3 population activity in Vgat-Het mice, to a greater extent than it did in WT mice. This suggests that groups of neurons with abnormally elevated correlations may contribute to the impaired behavioral performance of Vgat-Het mice. Furthermore, an increased number of L2/3 neurons in wS1 of Vgat-Het mice responded to the non-rewarded auditory tone, which likely contributes to premature behavioral reports. Collectively, we show that a decrease of SynGAP in inhibitory interneurons results in circuit dysfunction in the primary sensory cortex, characterized by elevated responses to irrelevant sensory stimuli and increased detrimental correlations.

## Results

### Whisker-guided tactile Go/No-go task in head-restrained mice

Although *SYNGAP1* expression was detected in inhibitory interneurons in the cerebral cortex, there has been no quantitative comparison of expression level across major neuronal subtypes. We analyzed *SYNGAP1* expression in cortical interneuron subtypes using a published web database (http://research-pub.gene.com/NeuronSubtypeTranscriptomes) that includes the transcriptome dataset collected by the Allen Brain Institute (Huntley et al., 2020; Tasic et al., 2016). *SYNGAP1* expression was detected in all three major inhibitory interneuron subtypes and excitatory cells in the cortex, consistent with previous studies (**Figure S1A**) (Zhang et al., 1999). A mouse line carrying ‘floxed’ *SYNGAP1* (*SYNGAP1^fl/fl^*) was previously reported (Ozkan et al., 2014). We generated inhibitory interneuron-specific *SYNGAP1* heterozygous (*Vgat^Cre/+^;SYNGAP1^fl/+^* or simply ‘Vgat-Het’) and wild-type (*Vgat^Cre/+^;SYNGAP1^+/+^*) mice by crossing *SYNGAP1^fl/+^* mice with Vgat-IRES-Cre (*Vgat^cre/cre^*) mice (**Figure S1B**). Inspired by prior work demonstrating altered tactile processing in the pan-neuronal *SYNGAP1* heterozygous knock-out (Llamosas et al., 2021; Michaelson et al., 2018), we first tested whether or not interneuron-specific *SYNGAP1* haploinsufficiency impacts tactile perception using a head-fixed whisker detection task (**Figure 1A**). Adult mice were water-restricted, habituated and then trained to report, by licking or withholding licking a reward port, whether facial whiskers received a brief sinusoidal deflection (25 Hz for 1 s, peak speed ~800 degrees s^−1^) (**Figure 1B**). Each trial began with a brief auditory tone (0.1 s, 8 kHz tone, 70 dB). This was immediately followed by 1.5 s ‘No-lick’ window, and licking during this period aborted the trial (**Figure 1C**). At 2 s after the offset of the auditory tone, the whisker deflection was delivered on 60 % of all trials (‘Go’ trials). The ‘response window’ was defined as 0.2-3.2 s after the onset of whisker deflection. Go trials resulted in a ‘hit’ when the mouse licked the water port within the response window and received a drop of water. In the remainder of trials (‘No-go’ trials; 40 %), whiskers were not deflected. Licking during the response window in the absence of whisker stimulus resulted in a 3 s time-out. The probability of correct choices during ‘Go’ (presence of whisker stimulus) and ‘No-go’ (absence of whisker stimulus) trials was monitored across training. Trial outcomes comprised a mixture of successful detection (‘hits’) and failed detection (‘misses’) following Go trials, as well as correct behavioral responses (‘correct rejections’) and incorrect responses (‘false alarms’) following No-go trials (**Figure 1D**).

**Figure 1.**
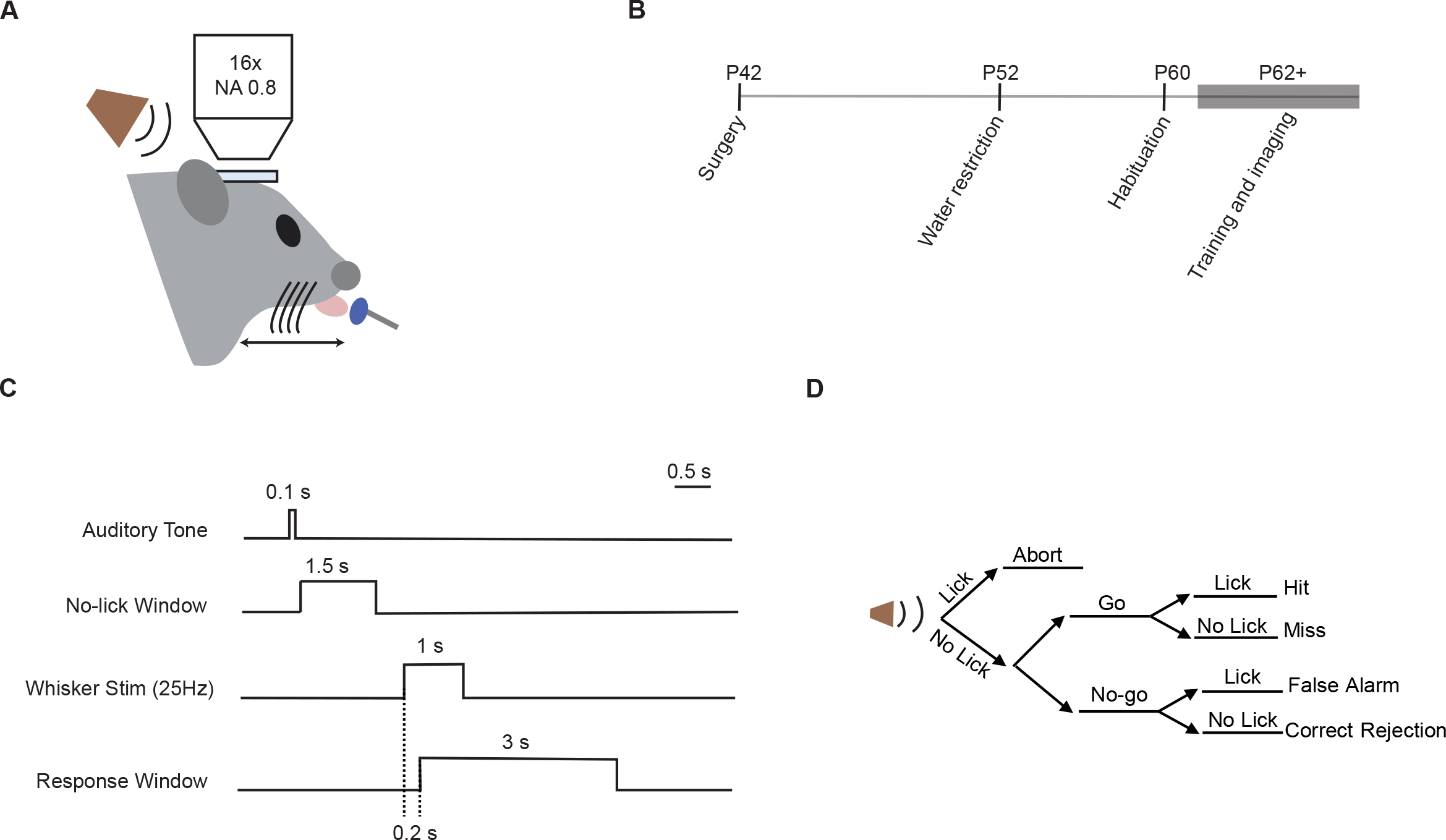
Tactile detection task in interneuron-specific *SYNGAP1* heterozygous mice. (A) Schematic showing the experimental set-up. Head-fixed mice were trained to lick a water port if facial whiskers were sinusoidally vibrated at 25 Hz with a piezoelectric stimulator or to withhold licking in the absence of whisker vibration. Neural responses in L2/3 of wS1 were monitored using two-photon imaging of genetically encoded calcium indicator jGCaMP6f or jGCaMP7f. (B) Experimental timeline. ‘P’, postnatal. (C) Each trial began with a brief auditory tone and was subsequently followed by 1.5 s ‘No-lick’ window; licking during this period resulted in trial abortion. On 60% of trials (‘Go’), the whiskers were then vibrated at 25 Hz for 1 s. On 40% of trials (‘No-go’), the whiskers were not vibrated. (D) Trial structure and outcomes.

### Heterozygous knock-out of SYNGAP1 in inhibitory interneurons impairs task learning

Wild-type (WT) littermate control mice (*Vgat^Cre/+^; SYNGAP1^+/+^*) steadily improved their behavioral performance and became ‘expert’ (discriminability index d’ > 2) at the whisker Go/No-go detection task after 6-7 daily sessions, whereas the performance of Vgat-Het mice (*Vgat^Cre/+^; SYNGAP1^fl/+^*) hovered between 1< d’ <2 (**Figure 2A**). Performances of WT and Vgat-Het mice were comparable on session 1 (pre-training), but WT mice performed significantly better than Vgat-Het mice on session 7 (post-training) (p = 0.014) (**Figure 2B**). The difference in post-training performance (session 7) was driven by higher false alarm rates of Vgat-Het mice, although they also displayed a slightly lower hit rate (hit rate: p = 0.174; false alarm rate: p = 0.013) (**Figures 2C, 2D, S2A and S2B**). Pan-neuronal *SYNGAP1* heterozygous mice (KO-Het) also showed elevated false alarm rates (p = 0.002) (**Figure S2B**) (Michaelson et al., 2018), but their hit rates were comparable to the WT (**Figure S2A**). The fraction of correct trials (hit and correct rejection) was significantly reduced in both Vgat-Het (p = 0.0004) and pan-neuronal KO-Het mice (p=0.009) (**Figure S2C**). These results demonstrate impaired sensory learning in Vgat-Het mice, which was driven by elevated behavioral responses in the absence of relevant sensory stimulus.

**Figure 2.**
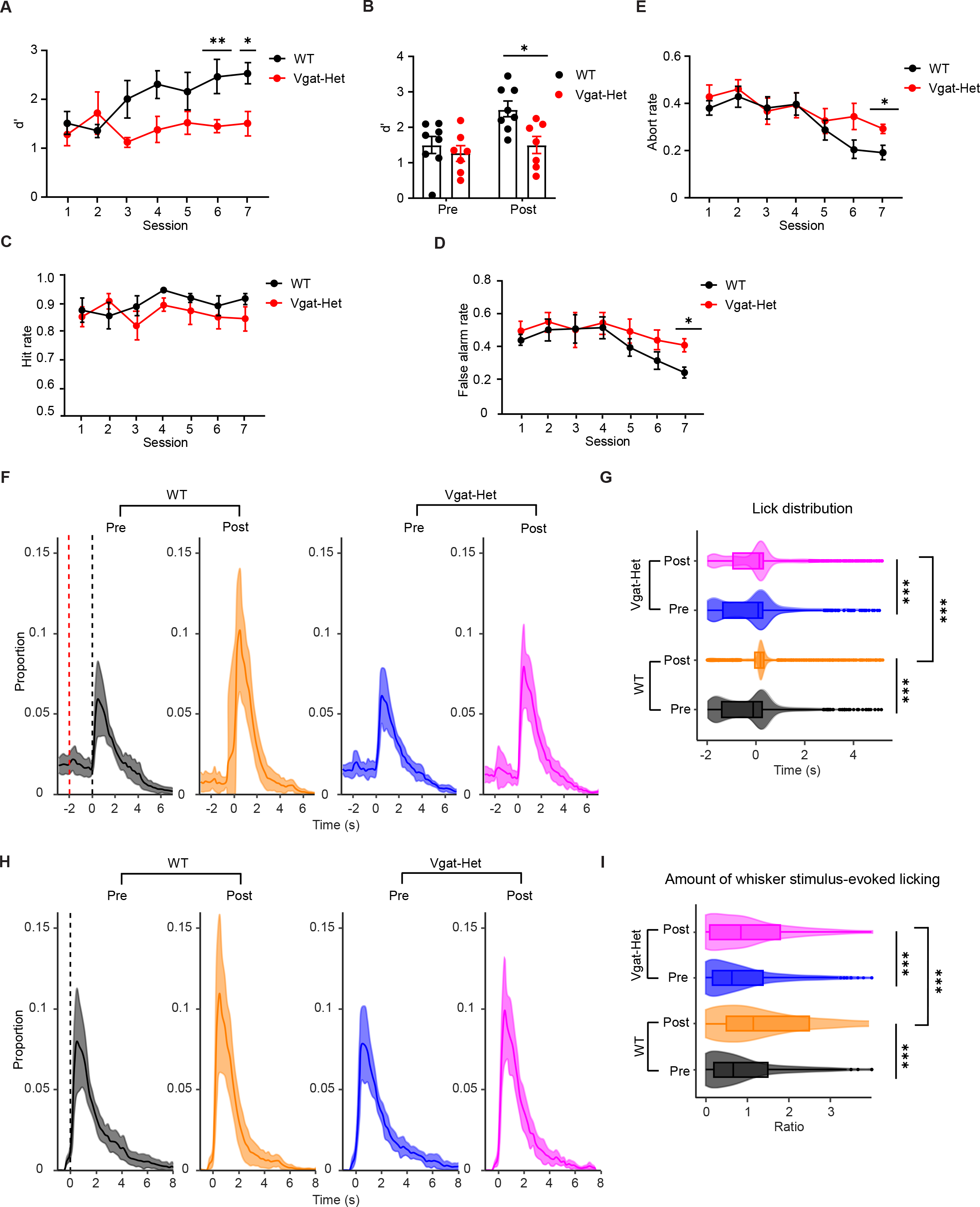
Vgat-Het mice show aberrant behavioral responses to sensory input. (A) Performance (d’) as a function of training sessions for WT (black) and Vgat-Het (red) animals. Error bars, mean ± S.E.M from 8 WT and 7 Vgat-Het mice. * p = 0.014, ** p = 0.004. (B) Performance (d’) of individual WT (black) and Vgat-Het (red) mice on session 1 (pre-training) and session 7 (post-training). Error bars, mean ± S.E.M from 8 WT and 7 Vgat-Het mice. (C) Hit rate for WT (black) and Vgat-Het (red) animals plotted over the course of training sessions. Error bars, mean ± S.E.M from 8 WT and 7 Vgat-Het mice. (D) False alarm rate for WT (black) and Vgat-Het (red) animals plotted over the course of training sessions. Error bars, mean ± S.E.M from 8 WT and 7 Vgat-Het mice. * p = 0.013. (E) Average fraction of aborted trials as a function of training sessions for WT (black) and Vgat-Het (red) animals. Error bars, mean ± S.E.M from 8 WT and 7 Vgat-Het mice. * p = 0.0205. (F) Average proportion of licks within 200 ms time bins, calculated as a fraction over the total number of licks combined across all trials including aborted ones. Red dashed line indicates the onset of auditory tone. Black dashed line indicates the expected onset of whisker stimulus. Error bars, mean ± S.E.M from 8 WT and 7 Vgat-Het mice. (G) Distribution of the first lick times during pre- and post-training sessions of WT versus Vgat-Het animals. Box plot indicates median and interquartile range. Well-trained WT mice (orange) make their first licks tightly around the expected onset time of whisker vibration, whereas a significant fraction of first licks made by post-training Vgat-Het mice (magenta) occurs before the expected onset of whisker stimulus. *** p < 0.0005. (H) Same as in panel *G* but excluding aborted trials. Black dashed line indicates the onset of whisker stimulus. Error bars, mean ± S.E.M from 8 WT and 7 Vgat-Het mice. (I) Fraction of licks that occur within 1 s after the expected onset of whisker stimulus. Box plot indicates median and interquartile range. WT animals (orange) target their licks more efficiently to the learned reward cue (within 1 s after the onset of whisker stimulus), compared to Vgat-Het mice (magenta). *** p < 0.0005.

Importantly, the fraction of aborted trials was elevated in Vgat-Het compared to WT mice in sessions 6-7, indicating an increased number of premature licks made during the ‘No-lick’ window (p = 0.0205) that precedes whisker stimulus onset (**Figure 2E**). The abort rate was also significantly higher in pan-neuronal *SYNGAP1* heterozygous mice compared to WT (p = 0.0210) (**Figure S2D**). The amount of water consumption (**Figure S2E**) or locomotor activity measured using open-field exploration (**Figure S2F**) was not altered in Vgat-Het mice (p = 0.090; p = 0.097). Therefore, the increased number of aborted trials cannot be accounted for by differences in motivation or general locomotor activity.

We monitored individual licks made by mice (**Figure S2G**) and compared their distribution across all trials in WT versus Vgat-Het mice before and after training for an equivalent number of sessions (**Figure 2F**). Compared to pre-training session, WT animals exhibited a decreased fraction of licks evoked by auditory tone in post-training session, whereas the number of licks made at the expected time of whisker stimulus onset increased (**Figure 2F**). Therefore, WT mice learned to effectively respond to reward-associated sensory input by shifting the time window of licking. Vgat-Het mice, on the other hand, continued to respond to the auditory tone in many trials even in session 7 (**Figure 2F**). We also compared distributions of time points when the first lick was made after auditory tone in each trial. First lick times, combined across all trials, were clustered around the expected time of whisker stimulus onset in expert WT mice, whereas they were more distributed in Vgat-Het mice (p < 0.0005) (**Figure 2G**). Next, we compared lick distribution in trials excluding aborted trials (**Figure 2H**). WT mice again showed more prominent training-induced increases in licking immediately after the expected time of whisker stimulus onset, compared with Vgat-Het mice (**Figure 2H**). We calculated the ratio between licks made within a 1 s time window after whisker stimulus onset versus licks outside this window (**Figure 2I**). Through training, both WT and Vgat-Het mice increased the number of licks within 1 s of whisker stimulus onset (p < 0.0005), but the change was significantly greater in WT mice (p < 0.0005) (**Figure 2I**). In summary, Vgat-Het mice show heightened behavioral responses to the irrelevant sensory input (auditory tone) and during the time window when the relevant stimulus (whisker vibration) is absent, compared to WT mice. Our results suggest that *SYNGAP1* haploinsufficiency in GABAergic neurons impairs learning by driving impulsive behavioral responses during a goal-directed tactile perception task.

### Neuronal sensitivity to whisker stimulus is slightly reduced

To characterize the relationship between behavioral performance and neural representations, we monitored responses of layer 2/3 neurons in the wS1 with two-photon calcium imaging of a genetically encoded calcium indicator (jGCaMP6f or 7f) as mice learned to perform the detection task (**Figures 3A and 3B**). We reasoned that, if *SYNGAP1* plays a role in the inhibitory circuit in the wS1, reducing its expression in cortical inhibitory interneurons should alter whisker input representation.

**Figure 3.**
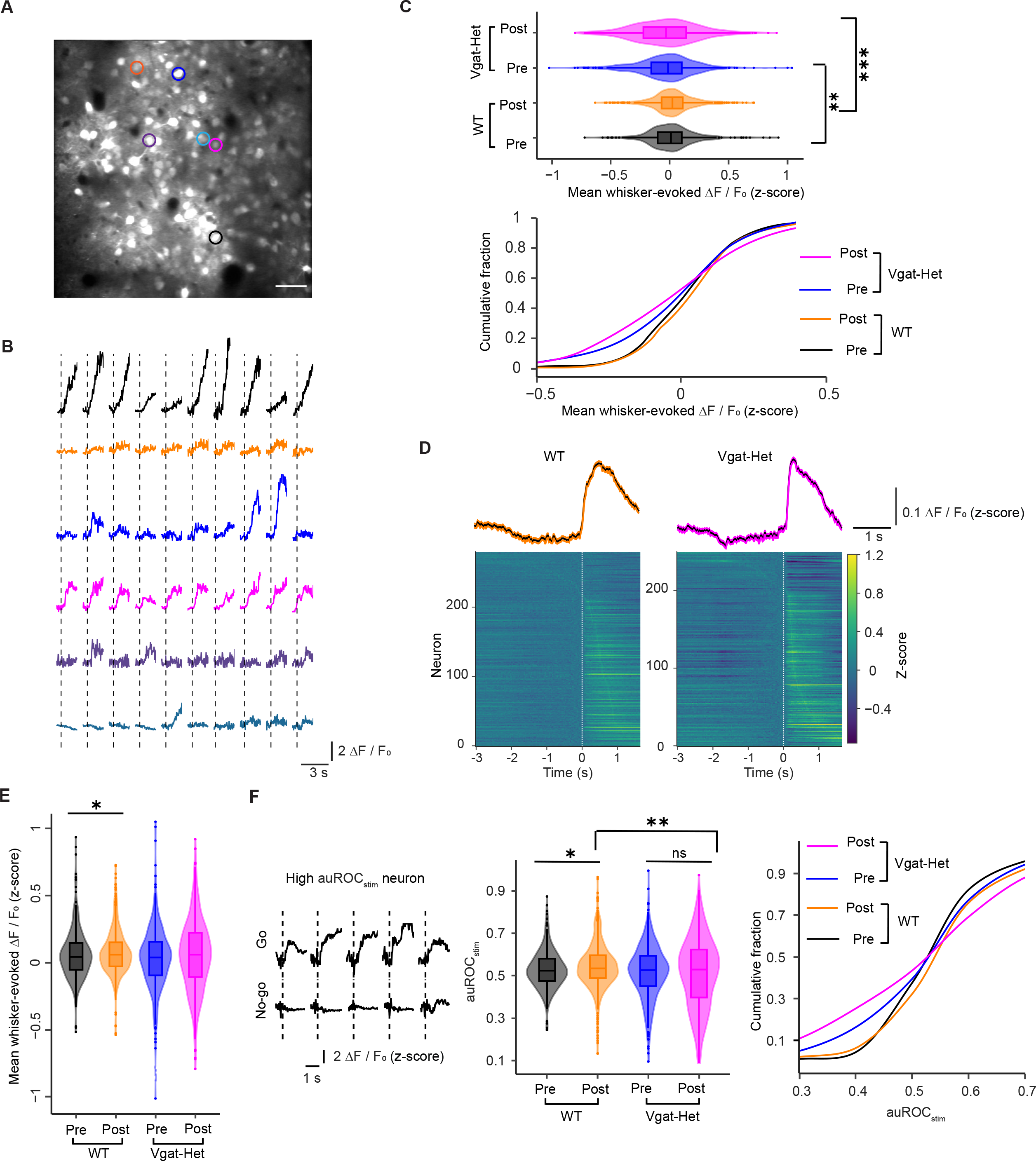
Whisker input representation by L2/3 neurons is impaired in Vgat-Het mice. (A) Example ‘field of view’ in the two-photon microscope, showing jGCaMP7f-expressing neurons. Scale bar, 100**μ**m. (B) Representative calcium transients evoked by whisker stimulation on Go trials. Different colors correspond to different cells as indicated by colored circles in *A*. Dashed lines indicate the onset of whisker deflection. (C) (Top) Magnitude of whisker-evoked responses in all cells on Go trials before and after training. Median [interquartile range, IQR] for WT-pre: 0.022 [−0.089, 0.116]; WT-post: 0.033 [− 0.056, 0.123]; Vgat-Het-pre: −0.004 [−0.142, 0.115]; Vgat-Het-post: −0.017 [−0.211, 0.153]. ** p = 0.002, *** p < 0.0005. (Bottom) Cumulative histograms of mean whisker-evoked *ΔF/F_o_*. (D) (Bottom) Each line corresponds to the average response of a single whisker-responsive neuron on Go trials in the post-training sessions. *ΔF/F_o_* traces were z-scored across all trials for each neuron and then averaged within Go trials. Neurons were pooled across animals and sorted by the time of peak activity. X-axis indicates time in relation to the onset of the whisker stimulus. (Top) Z-scored average *ΔF/F_o_* traces from post-training sessions in WT and Vgat-Het mice. Error bars, mean ± S.E.M from 5 WT (orange) and 5 Vgat-Het (magenta) mice. (E) Magnitude of whisker-evoked responses in whisker-responsive cells on Go trials. Median [IQR] for WT-pre: 0.045 [−0.051, 0.147]; WT-post: 0.060 [−0.026, 0.154]; Vgat-Het-pre: 0.039 [−0.096, 0.155]; Vgat-Het-post: 0.061 [−0.109, 0.221]. * p = 0.038. (F) (Left) Representative traces of a neuron with high auROC_stim_. (Middle) Area under receiver-operating-characteristic curve (auROC_stim_) quantifying how well an ideal observer could categorize the presence or absence of whisker stimulus (Go versus No-go trials) on the basis of the neural response. Median [IQR] for WT: auROC_stim_-pre = 0.524 [0.476, 0.580], auROC_stim_-post = 0.535 [0.487, 0.595]; Median [IQR] for Vgat-Het: auROC_stim_-pre = 0.526 [0.450, 0.593], auROC_stim_-post = 0.528 [0.397, 0.623]; * p = 0.013, ** p = 0.0049. (Right) Cumulative histograms of auROC_stim_.

We expressed jGCaMPs under pan-neuronal synapsin 1 promoter in the wS1 by virus injection and implanted a cranial window to enable optical access. We recorded activity from 613 neurons (5 WT mice) and 738 neurons (5 Vgat-Het mice) in the L2/3 of wS1. The average magnitude of neuronal response (*ΔF/F_o_*) evoked by whisker stimulation modestly decreased in Vgat-Het compared with WT mice (pre-training: p = 0.002; post-training: p < 0.0005) **(Figure 3C**). We found 41.0 ± 6.54 % of WT neurons and 35.3 ± 3.67 % of Vgat-Het neurons imaged in wS1 to be responsive to whisker stimulation, indicating a slight reduction in the pool of whisker-responsive L2/3 neurons in Vgat-Het mice. When the analysis was restricted to whisker-responsive cells, however, WT and Vgat-Het mice showed comparable magnitude and temporal dynamics of calcium transients (**Figure 3D**). Magnitudes of evoked *ΔF/F_0_* among whisker-responsive cells were similar between WT and Vgat-Het mice (pre-training: p = 0.038; post-training: p = 0.114) (**Figure 3E**).

To quantify neuronal sensitivity to whisker stimulus in a trial-by-trial manner, we calculated area-under receiver-operating-characteristinc curve (auROC_stim_), which captures how well an ideal observer could categorize sensory stimulus (in our case, presence or absence of whisker deflection) based on the neural response (Kwon et al., 2016; Yang et al., 2016). We used stimulus-evoked *ΔF/F_o_* as a decision variable for individual trials. auROC_stim_ significantly increased through training in WT mice (pre-vs post-training: p = 0.013), whereas it remained unchanged in Vgat-Het mice (p = 0.491) (**Figure 3F**). auROC_stim_ was also significantly greater in WT mice compared with Vgat-Het mice that went through an equivalent number of training sessions (p = 0.0049) (**Figure 3F**). Our results suggest that *SYNGAP1* haploinsufficiency in cortical interneurons causes subtle yet significant decreases in neuronal sensitivity to whisker stimulus in the L2/3 of wS1 during training.

### Vgat-specific SYNGAP1 knock-out disrupts population coding of whisker stimulus

Correlated trial-to-trial fluctuation in stimulus-evoked responses severely limits the amount of sensory information encoded by a neuronal population in the cortical network. The correlated co-variability between pairs of neurons or ‘noise correlation’ is usually a small positive number (range: 0.05-0.25) (Cohen and Kohn, 2011) and is positively related to synaptic connectivity (Ko et al., 2011). Altered pairwise noise correlations have been observed in several mouse models of neurodevelopmental disorders (Antoine et al., 2019; Banerjee et al., 2016; Lazaro et al., 2019). We calculated pairwise noise correlations among whisker-responsive neurons using their stimulus-evoked *ΔF/F_o_* on individual trials (Kwon et al., 2018; Kwon et al., 2016). Noise correlations decreased through learning in both WT and Vgat-Het mice (p < 0.0005 for both comparisons), consistent with the learning-associated improvement in sensory encoding (Ni et al., 2018) (**Figure 4A**). In trained mice, noise correlation was elevated (p = 0.0016) and more widely distributed in Vgat-Het compared with WT (**Figure 4A**). To compare the width of distribution between mice, we calculated the interquartile range (IQR) of noise correlations in individual mice and found that IQR was significantly larger in Vgat-Het mice (p = 0.015) (**Figure 4B**). We interpret this result as a greater number of positively and negatively correlated neuronal pairs being present in Vgat-Het mice compared to WT (Harris and Thiele, 2011).

**Figure 4.**
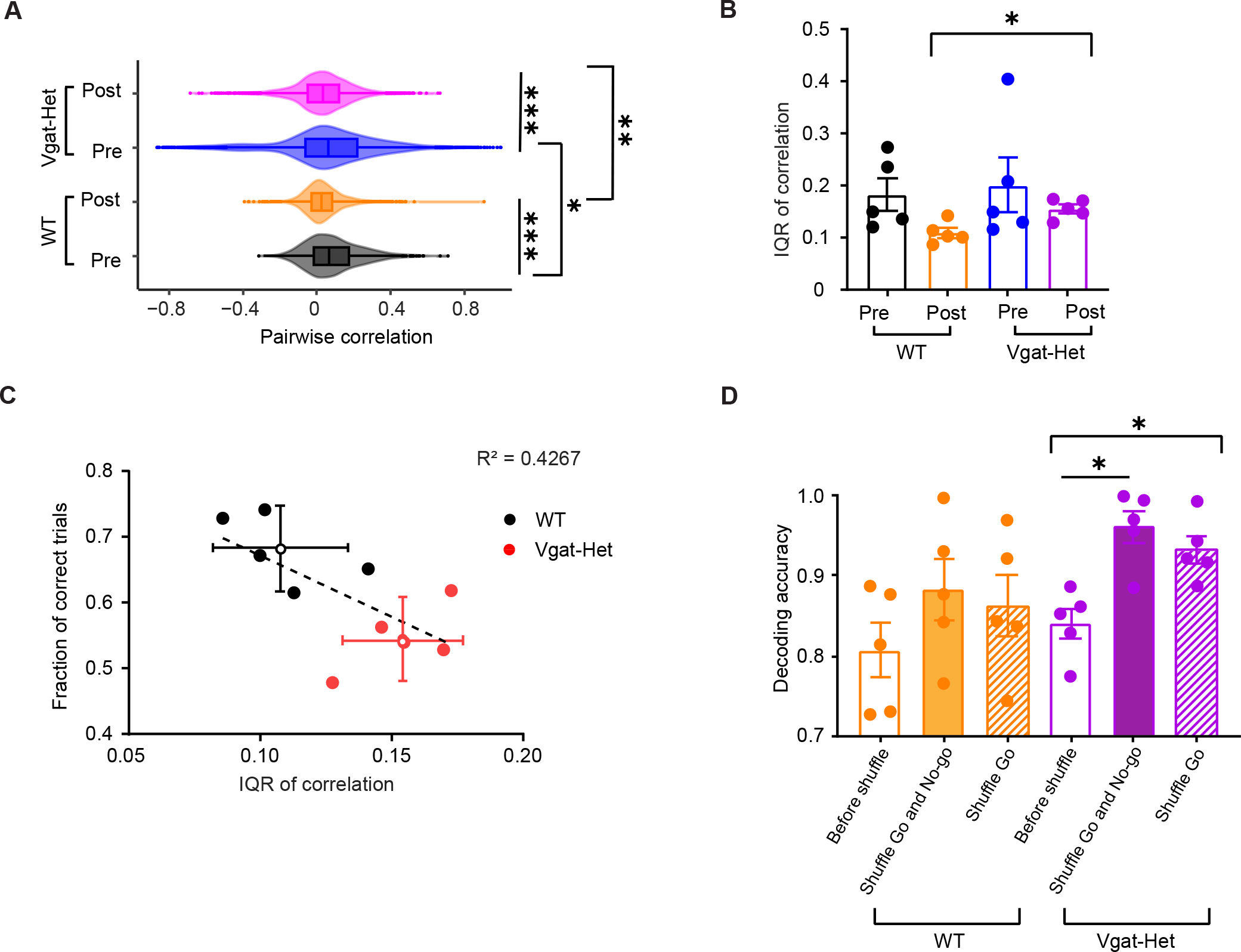
Noise correlation is disrupted in Vgat-Het mice. (A) Pairwise noise correlation calculated using all trials except the aborted ones. Median [IQR] for WT-pre: 0.067 [−0.013, 0.175]; WT-post: 0.023 [−0.027, 0.083]; Vgat-Het-pre: 0.063 [−0.064, 0.218]; Vgat-Het-post: 0.032 [−0.048, 0.120]. * p = 0.019, ** p = 0.0016, *** p < 0.0005. (B) IQR of distribution of noise correlations. Each dot represents a median IQR of a mouse. * p = 0.015. (C) Relationship between animal’s task performance and the distribution of pairwise noise correlation quantified using IQR. Dots indicate individual mice. Error bars, mean ± 95% C.I. p = 0.0407. Distance between WT and Vgat-Het centeroids: 0.144; 95% C.I. of null distribution [0.049, 0.053]. (D) Testing the impact of noise correlation on accuracy of decoding stimulus conditions from population activity. The support vector machine (SVM) classifier was trained to predict stimulus conditions on each trial based on the population activity. Bar graphs show average decoding accuracy before and after removing noise correlation by shuffling trial labels within both Go and No-go trials (solid) or within Go trials only (hashed). Each dot represents a mouse. Shuffle within Go and No-go: * p = 0.0121; Shuffle within Go: p = 0.0122, paired *t*-tests.

How does a wider distribution of noise correlations impact an animal’s behavior and/or encoding of sensory information in Vgat-Het mice? To begin to answer this question, we plotted IQR of noise correlations against performance for individual mice (*i.e*., fraction of correct trials) (**Figure 4C**). There was a negative relationship between distribution of noise correlations and task performance (R^2^ = 0.427); WT and Vgat-Het mice formed distinct clusters on this plot (**Figure 4C**). This suggests that the presence of aberrant noise correlations is likely to contribute to the impaired task performance in Vgat-Het mice. Next, we compared the amount of sensory information encoded by the L2/3 neuronal population. We decoded the stimulus condition (presence or absence of whisker deflection) using a support vector machine-based classifier trained on evoked *ΔF/F_o_* of the L2/3 neurons. The stimulus condition in individual trials could be predicted with about 80% accuracy in both WT and Vgat-Het mice (**Figure 4D**). We then removed noise correlations by shuffling trial labels in the same trial type (Go and No-go) and asked if the decoding of stimulus information could be improved. Removing noise correlations in both Go and No-go trials had mixed effects in WT mice with a slight improvement in decoding accuracy that did not meet statistical significance (Go and No-go: p = 0.3417; Go only: p = 0.475) (**Figure 4D**). For Vgat-Het mice, on the other hand, removing noise correlations in both Go and No-go trials or Go trials only significantly improved the decoding accuracy (Go and No-go: p = 0.0121; Go only: p = 0.0122) (**Figure 4D**). Therefore, a decrease of *SYNGAP1* expression in GABAergic interneurons introduces aberrant information-limiting noise correlation in the L2/3 population, which reduces the amount of sensory information.

### Vgat-specific SYNGAP1 knock-out increases cortical representation of irrelevant sensory input

Compared with WT mice, Vgat-Het mice make more licks in response to the auditory tone at the beginning of trials, resulting in an increased number of aborted trials (**Figure 2E**). We hypothesized that neuronal responses to the auditory tone might be abnormally elevated in Vgat-Het mice. To test this, we compared the activity of L2/3 wS1 neurons around the onset of the auditory tone. We focused on trials that do not contain licks immediately following the onset of tone presentation to exclude confounding effects of increased licking (**Figures 5A and 5B**). The fidelity of auditory tone-evoked responses was quantified as the fraction of trials in which statistically significant responses were elicited following the tone presentation. The auditory response fidelity was significantly elevated in Vgat-Het as compared to WT mice in both pre- and post-training sessions (p < 0.0005, Kolmogorov-Smirnov test) (**Figure 5C**). We then asked if auditory response fidelity is elevated in mice showing higher abort rates, by plotting the trial abort rate against average auditory response fidelity for individual mice. We found a positive correlation between these two metrics (R^2^ = 0.2284), although it was not statistically significant (**Figure 5D**). Analysis based on distance between centeroids of Vgat-Het and WT data points showed that they formed distinct clusters on this plot with Vgat-Het mice showing elevated auditory response fidelity and an increased number of aborted trials. We also compared the average magnitude of tone-evoked *ΔF/F_o_* between Vgat-Het and WT mice. The response magnitude was significantly greater in Vgat-Het as compared to WT mice in both pre- and post-training sessions (pre: p = 0.0014, post: p < 0.0005) (**Figure 5E)**. We conclude that a greater proportion of L2/3 neurons in Vgat-Het mice respond to the sensory input (tone) unrelated to reward acquisition, which might contribute to the elevated abort rate.

**Figure 5.**
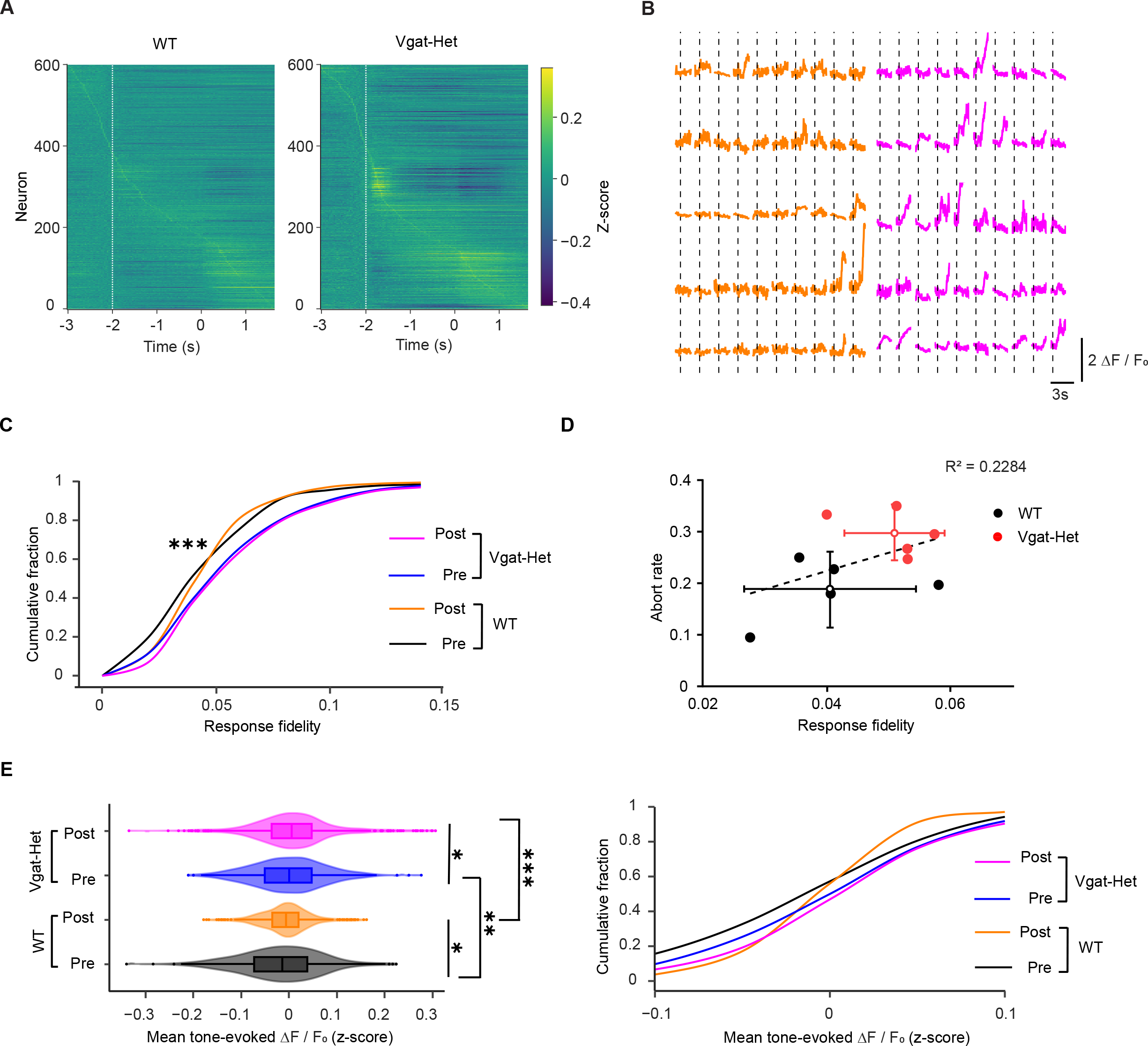
L2/3 neurons in Vgat-Het mice show enhanced responses to sensory input unrelated to the reward acquisition. (A) Each line corresponds to a single neuron. *ΔF/F_o_* traces were z-scored first across all trials for each neuron and then averaged within non-aborted trials. X-axis indicates time in relation to the onset of whisker stimulation. White dashed line indicates the onset of auditory tone. (B) Representative fluorescence traces of auditory tone-responsive neurons. WT (orange) and Vgat-Het (magenta). (C) Cumulative density plots showing fidelity of tone-evoked responses in the L2/3 neurons of wS1 before and after training. *** p < 0.0005, Kolmogorov-Smirnov test. (D) Relationship between abort rate and fidelity of tone-evoked responses. p = 0.1624. Distance between WT and Vgat-Het centeroids: 0.109; 95% C.I. of null distribution: [0.038, 0.042]. (E) (Left) Mean tone-evoked responses in all cells of non-aborted trials. Median [IQR] for WT-pre: −0.013 [−0.073, 0.037];WT-post: −0.006 [−0.035, 0.019], * p = 0.034; Vgat-Het-pre: 0.00 [−0.050, 0.046]; Vgat-Het-post: 0.005 [−0.036, 0.047], * p = 0.029. ** p = 0.0014, *** p < 0.0005. (Right) Cumulative histograms of mean tone-evoked *ΔF/F_o_*.

## Discussion

Previous studies demonstrated the role of SynGAP in regulating the development and plasticity of excitatory neurons in the neocortex (Clement et al., 2012; Ozkan et al., 2014). More recently, it has been reported that SynGAP also controls the migration and connectivity of cortical inhibitory interneurons (Berryer et al., 2016; Su et al., 2019; Sullivan et al., 2020). In the present study, we tested if and to what extent *SYNGAP1* haploinsufficiency in GABAergic cells contributes to cognitive and cortical circuit abnormalities.

Experiments in head-fixed mice performing whisker-guided sensory detection tasks are widely used for probing circuit dysfunction in autism models including pan-neuronal *SYNGAP1* and interneuron-specific *SHANK3* knock-out mice (Chen et al., 2020; Michaelson et al., 2018). Head-fixed preparations were also adopted in studies examining visual processing in autism models (Batista-Brito et al., 2017; Del Rosario et al., 2021; Goel et al., 2018). In most previous studies, however, neural activity and task performance were not measured simultaneously (except see (Del Rosario et al., 2021)). By combining two-photon calcium imaging and quantitative behavioral tasks in the same animal, we directly tested the relationships between neural activity and behavior in GABAergic *SYNGAP1* haploinsufficiency.

Knocking out a copy of *SYNGAP1* in Vgat +ve inhibitory interneurons was sufficient to cause learning deficits characterized by impaired detection task performance and increased tendency to respond in the absence of relevant sensory input (**Figure 2**). We also confirmed that pan-neuronal *SYNGAP1* knock-out impaired the task performance and elevated the false alarm rate as previously reported (Michaelson et al., 2018). An increased false alarm rate was also observed in models of other neurodevelopmental disorders, including *Fmr1* knock-out mice (Goel et al., 2018). A novel finding of this study is that both pan-neuronal and Vgat-specific *SYNGAP1* knock-out mice show an increased number of premature responses to the sensory input unrelated to reward acquisition (**Figure S2**). Based on these results, we conclude that *SYNGAP1* expression in inhibitory interneurons is required for generating appropriate behavioral responses to sensory input during goal-directed behaviors.

Behavioral deficits described above were accompanied by a disrupted representation of whisker input in the L2/3 of wS1, characterized by the following: (i) reduced neuronal sensitivity to whisker stimulus (auROC_stim_) (**Figure 3**), (ii) increased amount of information-limiting correlations (**Figure 4**), and (iii) elevated responses to sensory input unrelated to reward acquisition such as the auditory tone (**Figure 5**). Noise correlations have been previously measured in several mouse models of neurodevelopmental disorders. Mean noise correlations were found to be decreased in *Fmr1* KO (Antoine et al., 2019), decreased in PV neuron-specific *MeCP2* KO, but increased in SST-specific MeCP2 KO (Banerjee et al., 2016), and unchanged in wS1 of *CNTNAP2* KO (Antoine et al., 2019), but altered in prefrontal cortex differently depending on neuronal subtypes (Lazaro et al., 2019). However, until now it has been unclear to what extent altered noise correlations contribute to altered task performance in these mouse models. We report that noise correlations in inhibitory interneurons in primary somatosensory cortex are distributed with greater variance and that the increased width of the distribution predicts poorer task performance in Vgat-Het mice. Removing noise correlations substantially improved sensory information, consistent with a greater amount of detrimental correlations in Vgat-Het mice. Our findings highlight the altered distribution of noise correlations as a potential circuit endophenotype that could be utilized to stratify different neurodevelopmental disorders.

While we report cortical circuit disruptions associated with specific observed behavioral deficits, we do not claim that wS1 is the only brain structure contributing to the deficits described here. *SYNGAP1* is most abundantly expressed in the cerebral cortex and hippocampus during brain development, but a modest level of expression is also detected in striatum (Araki et al., 2020; Komiyama et al., 2002). Although *Vgat* is expressed in both GABAergic and glycinergic neurons (Wang et al., 2009), glycinergic neurons are sparse in the neocortex where *SYNGAP1* is abundant. Therefore, our intersectional genetic approach predominantly targets *SYNGAP1* expression in GABAergic inhibitory interneurons in cortex and hippocampus. The behavioral deficits reported here are likely to originate from circuit disruptions in these areas. Future studies could examine the role of *SYNGAP1* in striatum and the associated behaviors, using circuit-specific manipulations. Another potential caveat in interpreting our results is that *SYNGAP1* haploinsufficiency may disrupt the ascending sensory processing pathway. This is unlikely, however, as *SYNGAP1* haploinsufficiency has little effect on the development of the somatosensory barrel map or thalamocortical innervation in wS1 (Barnett et al., 2006).

Whether circuit abnormalities observed in the wS1 causally drive the observed behavioral deficits of Vgat-Het mice, or if other cortical regions are also involved in this process, are important remaining questions. *SYNGAP1* haploinsufficiency exerts differential effects on PV neurons - a major cortical GABAergic interneuron subtype - in different cortical regions. PV neurons are reduced in number in the prefrontal cortex but not in wS1 of pan-neuronal *SYNGAP1* heterozygous knock-out mice (Sullivan et al., 2020). Interestingly, in these mice expression of the AMPA receptor subunit GluA2 is selectively elevated in PV neurons that are located in wS1 but not those in the prefrontal cortex (Sullivan et al., 2020). *SYNGAP1* expression in PV cells is known to be important for perisomatic innervation of excitatory pyramidal neurons in wS1 (Berryer et al., 2016). *SYNGAP1* is also expressed in SST and VIP inhibitory interneurons (**Figure S1A**). Neuronal subtype-specific functions of SynGAP and how these are disrupted by *SYNGAP1* haploinsufficiency warrants further investigation.

Collectively, our results add to growing evidence highlighting the contribution of cortical inhibitory interneurons to sensorimotor abnormalities in neurodevelopmental disorders (Contractor et al., 2021). Furthermore, we identify circuit-level endophenotypes in the primary somatosensory cortex that likely contribute to sensorimotor impairments. These findings have direct implications for studies focusing on circuit dysfunction in autism spectrum disorders.

---

**Supplemental Figure 1.**
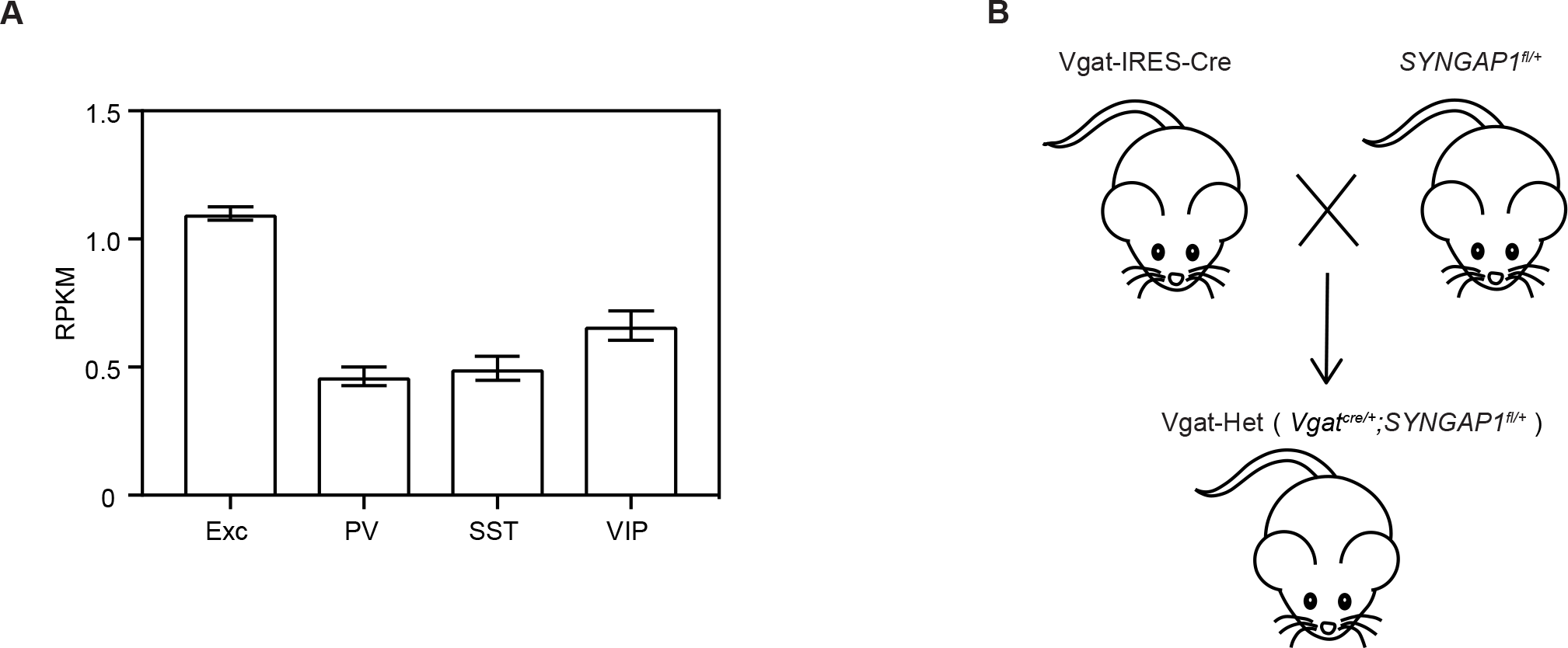
(A) Expression of *SYNGAP1* transcripts in excitatory (Exc), and inhibitory – parvalbumin (PV), somatostatin (SST) and vasointestinal peptide (VIP) -- neurons based on an analysis of a single-cell transcriptomics dataset. Transcript level was quantified as reads per kilobase of transcript, per million mapped reads (RPKM). (B) Breeding scheme for knocking out a copy of *SYNGAP1* in inhibitory interneurons. *SYNGAP1*^fl/+^ mouse was crossed with Vgat-IRES-Cre to create *Vgat^Cre/+;^ SYNGAP1^fl/+^* mice (Vgat-Het). *Vgat^Cre/+;^ SYNGAP1^+/+^* mice were used as wild-type control (WT).

**Supplemental Figure 2.**
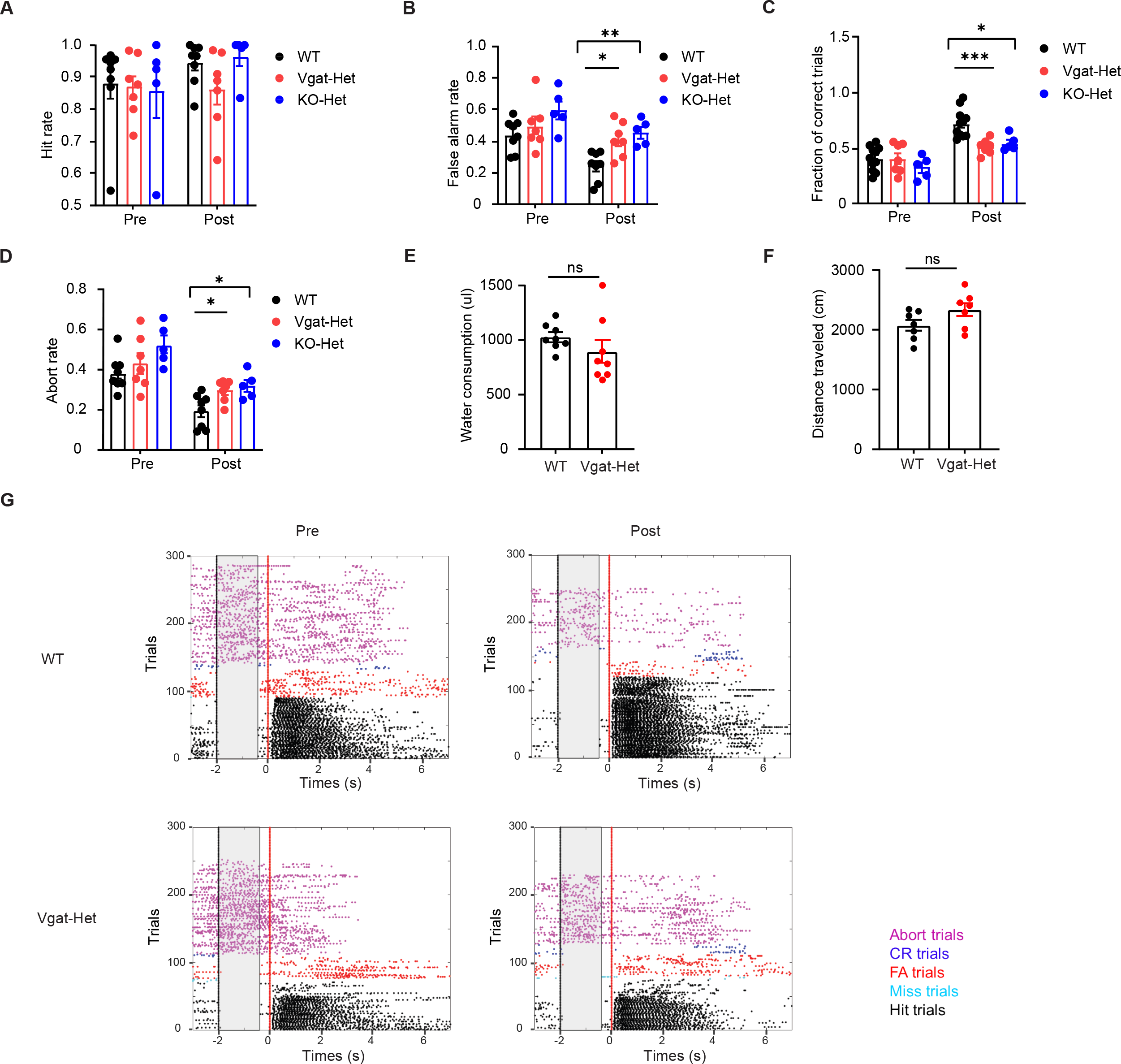
(A) Hit rate of individual WT (black; n = 8), Vgat-Het (red; n = 7) and KO-Het (blue; n = 5) mice on session 1 (pre-training) and session 7 (post-training). Error bars, mean ± S.E.M. Comparison of post-training sessions: p=0.174 (WT against Vgat-Het), p=0.179 (WT against KO-Het). (B) False alarm rate of individual WT (black; n = 8), Vgat-Het (red; n = 7) and KO-Het (blue; n = 5) mice on session 1 (pre-training) and session 7 (post-training). Error bars, mean ± S.E.M. *p=0.013, **p=0.002. (C) Fraction of correct trials on pre-learning and post-learning sessions, calculated as (Hits + Correct Rejection) / all trials including aborted ones. Error bars, mean ± S.E.M from 8 WT, 7 Vgat-Het mice and 5 SynGAP KO-Het mice. * p=0.009, *** p = 0.0004. (D) Abort rate on pre-learning and post-learning sessions, calculated as (Hits + Correct Rejection) / all trials including aborted ones. Error bars, mean ± S.E.M from WT (black; n = 8), Vgat-Het (red; n = 7) and KO-Het (blue; n = 5). *p=0.0205 (WT against Vgat-Het), *p=0.0210 (WT against KO-Het). (E) Water consumption of each day. p=0.090. (F) Open field test distance traveled within 8 minutes. p=0.097. (G) Example sessions from WT and Vgat-Het animals, before versus after training in the detection task (session 1 versus session 7), showing trials sorted by type. Ticks indicate lick times and are colored by five trial outcomes; hit (black), miss (cyan), false alarm (red), correct rejection (blue) and aborted (magenta) trials.

## Methods

### Mice

All procedures were in accordance with protocols approved by the University of Michigan Animal Care and Use Committee. We report data from simultaneous calcium imaging and behavior from 5 *SYNGAP1* WT mice (*Vgat^Cre/+^; SynGAP^+/+^*) and 5 Vgat-Het mice (*Vgat^Cre/+^; SynGAP^fl/+^*) (Jackson labs) with a C57BL/J6 background, with ages ranging from 8 to 15 weeks. For results describing behavioral phenotypes (**Figure 2**), we used 8 *SYNGAP1* WT mice (*Vgat^Cre/+^; SynGAP^+/+^*; 5 of 8 with calcium imaging), 5 pan-neuronal *SYNGAP1* heterozygous mice (*Vgat^+/+^; SynGAP^fl-stop/+^*) and 7 GABA-specific heterozygous mice (*Vgat^Cre/+^; SynGAP^fl/+^*; 5 of 7 with calcium imaging) (Jackson labs) with a C57BL/J6 background, with ages ranging from 8 to 15 weeks. Both sexes were used. Mice were housed in a vivarium with a reversed light-dark cycle (12 h each phase). Experiments occurred during the dark phase. After recovery from head-post surgery (see below), mice were singly housed and water-restricted by giving them 1 mL per day. Mouse weight did not decrease below 70% of the starting weight.

### Behavioral Task

Head-restrained mice were trained to perform a Go/No-go whisker detection task using a behavioral apparatus controlled by BPod (Sanworks). Mice were placed in an acrylic (4.5 cm inner diameter) tube. For 7–10 d before training, mice received 1 ml per day of water. Mice were weighed prior to and after training sessions to ensure the amount of water consumed. In the first 3 sessions (‘Habituation’), mice were allowed to freely lick at the water port positioned near their snout. Each time the tongue crossed the infrared beam to touch the water port, the mouse received a drop of water (~7 μL). For training in ‘Go/No-go’ sessions (7-10 days), facial whiskers were threaded through a plastic mesh attached to a piezoelectric actuator (CTS) and were deflected for 1 s with sinusoidal deflection (rostral to caudal) at 25 Hz on Go trials (60% of trials). On No-go trials (40% of trials), the whiskers were not deflected. Starting 1 s after trials started, a 0.1 s auditory tone (8 kHz tone, ~70 dB SPL) was delivered, followed by a 1.5 s ‘No-lick’ window. If mice lick during this window, i.e. premature licking, the trial was aborted. Licks occurring during the first 0.2 sec after the onset of whisker deflection had no consequence. The ‘response window’ was defined as 0.2-3.2 s after the onset of whisker deflection. Go trials resulted in a ‘hit’ when the mouse licked the water port within the response window and received a drop of water. A ‘miss’ occurred if mice did not lick within the response window, and no reward or punishment was delivered. The No-go trials resulted in a ‘false alarm’ if mice licked within the response window, and mice were punished by 3 s time-out. Licking during time-out resulted in an additional time-out. A ‘correct rejection’ occurred if mice did not lick within the response window on No-go trials. During all sessions, ambient white noise (cut off at 40 kHz, ~60 dB SPL) was played through a separate speaker to mask any other potential auditory tones associated with the movement of the piezoelectric actuator. No more than 3 trials of the same type occurred in a row. The fraction of correct trials (Fraction Correct) was defined as the number of hit and correct rejection trials divided by the total number of trials. The hit rate was defined as the number of hits divided by the number of Go trials. The false alarm rate was defined as the number of false alarms divided by the number of No-go trials. The abort rate was defined as the number of aborted trials divided by the number of all trials.

### Surgery and Virus Injection

Mice were anesthetized with 1% isoflurane throughout surgery and kept on a thermal blanket to maintain body temperature. The scalp and periosteum over the skull were carefully removed. A circular craniotomy was made on the left hemisphere (3.0 mm diameter) with the dura left intact. The center of the craniotomy was located over the wS1 barrel cortex (3.5 mm lateral and 1.3 mm caudal relative to Bregma). Injections were performed unilaterally using a beveled glass pipette (30-50 μm diameter) mounted on an oil-based hydraulic micromanipulator (Narishige). Adeno-associated virus for expressing jGCaMP6f or jGCaMP7f under the synapsin-1 promoter (AAV1-syn-jjGCaMP6f-WPRE-SV40, Addgene, 100837; AAV1-syn-jjGCaMP7f-WPRE, Addgene, 104488) was injected into the wS1 at depth of 250 μm below the dura and at a rate of 1 nL/sec (100 nL total). Both WT and Vgat-Het groups contained three jGCaMP6f- and two jGCaMP7f-injected mice. The injection was made at 3 different locations on the cortical surface around the coordinates given above. The craniotomy was covered with a glass window after the injection. The window was made by gluing two pieces of coverslip glass together. The smaller piece (3.0 mm diameter) was placed into the craniotomy and while the larger piece (4.0 mm diameter) was glued to the bone surrounding the craniotomy. Cyanoacrylate adhesive (KrazyGlue) and dental acrylic (Jet Repair Acrylic) were used to secure a titanium head post in place on the skull. Silicone elastomer (Kwik-Cast, WPI) was placed over the window for protection during the recovery period. The mouse was allowed to recover from surgery for at least 10 days before moving to water restriction. Imaging started 3-5 weeks after surgery.

### Two-Photon Calcium Imaging of Layer 2/3 neurons

Images were acquired on a Scientifica two-photon microscope (Hyperscope) equipped with an 8 kHz resonant scanning module, 2 GaAsP photomultiplier tube modules, and a 16× 0.8 NA microscope objective (Nikon). jGCaMP was excited at 960 nm (40-60 mW at specimen) with an InSight X3 tunable ultrafast Ti:Sapphire laser (Spectra-Physics, Santa Clara, CA, USA). Imaging fields were restricted to areas where jGCaMP expression overlapped with the center of the cranial window (3.5 mm lateral and 1.3 mm caudal to Bregma). The beam was focused to 150 – 250 μm from the cortical surface. The field of view ranged from 458 μm × 344 μm to 275 μm × 207 μm. Images were acquired with a resolution of 512 × 512 pixels at 30 Hz using ScanImage. A movie for a single trial consisted of 140 frames.

### Image Analysis

Image stacks were processed using Suite2P pipeline (Stringer and Pachitariu, 2019). After correcting for motion, regions of interest (ROIs) were selected and then manually curated to remove ROIs that were not neurons. The neuropil fluorescence time series was multiplied with a correction factor of 0.7 and then subtracted from the raw fluorescence time series to obtain the corrected fluorescence time series: *F_corrected_(t) = F_raw_ − F_neuropil_* * 0.7. *ΔF/F_o_* was calculated as (*F* − *F_o_*) / *F_o_*, where *F_o_* represents the baseline fluorescence calculated by determining the average fluorescence (*F*) in the preceding 8 frames time window from whisker stimulus onset. Evoked *ΔF/F_o_* responses were calculated as the average *ΔF/F_o_* over the 10 frames following the expected whisker stimulus onset time.

### Single neuron analysis

To quantify the response fidelity, we calculated the percentage of whisker stimulus-responsive trials for each neuron. If 15 frames (0.5 s) following the onset of whisker stimulus contain 3 or more frames with fluorescence intensity > (mean over 8 frames preceding the stimulus onset time + 3 x standard deviations) or < (mean over 8 frames preceding the stimulus onset time − 3 x standard deviations), the trial was considered responsive. To assign each neuron as ‘responsive’ or ‘unresponsive’, we used 0.02 as the response fidelity cutoff so that neurons with response fidelity > 0.02 were considered ‘responsive’. The receiver operating characteristic (ROC) analysis was used to calculate auROC_stim_ (Kwon et al., 2016). We used across-trial z-scores for the evoked *ΔF/F_o_* as a decision variable for each neuron. Trials were grouped by stimulus condition (present versus absent). The ROC curve was computed by systematically varying the criterion value across the full range of decision variable (using Python sklearn ‘roc_auc_score’ function). The area under the ROC curve (auROC) represents the performance of an ideal observer in categorizing trials based on the decision variable.

### Noise correlation analyses

We calculated across-neuron pairwise noise correlations between neuron pairs recorded at the same time in a single session, across trials sharing the same stimulus condition (presence or absence of whisker stimulus) (Kwon et al., 2018; Kwon et al., 2016). For each neuron, evoked *ΔF/F_o_* responses were first z-scored in Go and No-go trials separately. Pairwise noise correlation was then calculated as the Pearson correlation coefficient between vectors of concatenated z-scored responses for each pair of neurons.

### Population decoding analysis

Support Vector Machine classifier (SVM, Python sklearn package) was trained to discriminate the stimulus condition (Go versus No-go trials) based on a vector of z-scored evoked *ΔF/F_o_* for all responsive cells in each session. 5-fold cross-validation was performed by using a random 80% of trials for training and the remaining 20% for testing the classifier performance. Population decoding accuracy was calculated as the percentage of trials in the test set that was accurately predicted by the classifier. This was done 100 times and the average accuracy was used for comparison between animals. Go (60%) and No-go (40%) trials were used and randomly assigned to train and test subgroups. To evaluate how noise correlations contribute to neuronal performance, we randomly shuffled trial labels within the same trial type (Go or No-go) in neurons independently from each other by using numpy.random.permutation to remove correlated trial-by-trial variability. Then we used SVM as above to predict stimulus conditions based on the population activity and calculated the average accuracy.

### Open field test

Mice were habituated in the open field apparatus (30 × 30 × 30 cm) for 5 mins and then total distance traveled (cm) within 8 mins was recorded and analyzed using a software (ToxTrac). Each mouse was tested for 2 or 3 trials, and the distance was averaged across trials.

### Statistical tests

Statistical significance of differences between WT and Vgat-Het mice was assessed using unpaired Wilcoxon rank-sum tests unless mentioned otherwise. To test if WT and Vgat-Het mice form statistically distinct clusters on a scatter plot (**Figure 5D**), we calculated distance between two centeroids, each representing mean of data points from WT or Vgat-Het mice. We then derived a distribution representing ‘null hypothesis’ by shuffling labels (WT or Vgat-Het) associated with data points 100 times and calculating distance between centeroids each time. If the inter-centeroid distance is greater than central 95% of the ‘null’ distribution, WT and Vgat-Het were considered to be separate from each other.

## Resource availability

Original data reported in this paper will be shared by the lead contact upon request. This paper does not report original code. Source data and scripts used for data analysis will be deposited and publicly available.

## Acknowledgments

We thank members of the Kwon laboratory, Sara Aton and Pamela Raymond for critical reading of the manuscript. This work was supported by a Bridge to Independence grant from the Simons Foundation Autism Research Initiative (S.E.K).

## Author contributions

MZ and SEK designed experiments. MZ performed all experiments and analyzed data. MZ and SEK wrote the manuscript.

